# Enhanced epigenetic modulation via mRNA-encapsulated lipid nanoparticles enables targeted anti-inflammatory control

**DOI:** 10.1101/2025.02.24.639996

**Authors:** Tahere Mokhtari, Mohammad N. Taheri, Sarah Akhlaghi, Armin Aryannejad, Yuda Xiang, Vineet Mahajan, Kamyar Keshavarz, Amirreza Kiani, Samuel LoPresti, Ryan LeGraw, Kathryn A. Whitehead, Samira Kiani

## Abstract

Temporal transcriptional modulation of immune-related genes offers powerful therapeutic potential for treating inflammatory diseases. Here, we introduce an enhanced zinc finger (ZF)-based transcriptional repressor delivered via lipid nanoparticles for controlling immune signaling pathways *in vivo*. By targeting Myd88, an essential adaptor molecule involved in immunity, our system demonstrates therapeutic efficacy against septicemia in C57BL/6J mice and improves repeated AAV administration by reducing antibody responses. This epigenetic engineering approach provides a platform for safe and efficient immunomodulation applicable across diseases caused by imbalanced inflammatory responses.

## Introduction

Overactive immune responses pose significant challenges in various inflammatory diseases, often leading to severe tissue damage, organ dysfunction, and mortality^1^. This phenomenon, termed “cytokine storm”, manifests through uncontrolled release of pro-inflammatory mediators and has been implicated in conditions ranging from acute respiratory distress syndrome (ARDS) to sepsis ^2,3^. Current therapeutic approaches rely largely on broad-spectrum immunosuppressants, which can leave patients vulnerable to opportunistic infections and lead to significant adverse effects^4–7^.

Beyond acute inflammatory conditions, immune responses hamper the success of gene therapies, particularly for those exploiting adeno-associated virus (AAV) delivery^8^. Pre-existing humoral immunity against AAV excludes many patients from trials and compromises treatment outcomes^9,10^. Several factors influence viral vector immunogenicity, including capsid serotype, vector DNA, target tissue inflammatory state, and host immunity status^11,12^. While various strategies attempt to regulate immune responses against AAV vectors, existing approaches face limitations in overcoming pre-existing immunity and enabling vector redosing^13–16^.

Myd88, an essential adaptor for Toll-like receptors (TLRs) and upstream regulator of NF-κB signaling, represents a promising target for addressing both hyperinflammatory conditions and gene therapy challenges^17^. MyD88 signaling has been directly implicated in forming immune responses against AAV capsid, which leads to loss of transgene expression and limits effective administration of the virus^18^. Furthermore, MyD88 signaling is essential for survival against viral infections, making it an attractive target for modulating both innate and adaptive immunity to viral vectors^19^. Traditional approaches for targeting *Myd88*, such as small molecule inhibitors, have been explored but suffer from lack of specificity and proper tissue penetrance *in vivo*^20,21^.

While our previous work demonstrated that CRISPR-based epigenetic downregulation of *Myd88* delivered via AAVs can effectively dampen immune responses^22^, this approach faces limitations including potential immunogenicity of CRISPR and AAV components, lack of transient control, and challenges in scalability. To address these challenges, we introduce a novel method for synthetic immunomodulation using mRNA-based zinc finger transcriptional repression targeting *Myd88*. Our strategy involves zinc finger directed epigenetic modulation, allowing precise and reversible control over gene expression. Zinc finger proteins offer several advantages over other epigenetic-editing technologies, including low immunogenicity, smaller size for easier packaging and delivery, and high DNA binding specificity^23,24^. To avoid complexities associated with viral vehicles, we deliver the zinc finger-based epigenetic repressor mRNA via lipid nanoparticles (LNPs), providing an attractive platform for achieving transient gene regulation while offering potential for scale-up^25^.

## Results and Discussion

### Zinc finger-mediated repression with HP1a-KRAB efficiently downregulates *Myd88 in vitro*

In our previous studies, we established CRISPR-based epigenetic repressors using two methods: the direct fusion of modulatory elements to a catalytically inactive Cas9, and an indirect approach utilizing aptamers to guide MS2-fused transcriptional repressors to the CRISPR complex^22,26^. Building upon our prior findings, we sought to develop a zinc finger (ZF)-tethered Myd88-targeting repressor with enhanced efficacy and specificity. We designed a panel of 16 zinc finger sequences targeting various regions of the *Myd88* promoter and evaluated their repressive capacity when fused to HP1a-KRAB. Through systematic screening in mouse neuroblastoma (N2A) cells and quantitative real-time polymerase chain reaction (qRT-PCR), we identified zinc finger-effector 11 (ZFR11) as the most potent repressor of *Myd88* expression (Supplementary Fig. 1A, B).

We compared the functionality of ZFR11 to our previously optimized CRISPR-based system, which utilizes a 14-nucleotide truncated guide RNA, Cas9 nuclease, and MS2-HP1a-KRAB. qRT-PCR analysis revealed that ZFR11 achieved *Myd88* repression comparable to that achieved with an enhanced aptamer-mediated CRISPR repressor (Supplementary Fig. 1C). These findings reveal the *in vitro* functionality of the ZFR11 and underscore its potential as a highly efficient tool for modulating *Myd88* expression, offering an alternative to CRISPR-based approaches with enhanced safety profiles and greater flexibility in delivery methods.

### 306O10 LNP-mediated delivery enables effective transfection of diverse immune cell populations *in vitro* and *in vivo*

To enhance the efficiency of *Myd88* repression and enable transient immunomodulation, we constructed epigenetic modulators in form of mRNA (*in vitro-*transcribed), which were subsequently delivered into diverse cells via LNPs. LNP-mediated mRNA delivery provides several advantages over viral and other non-viral deliveries—most importantly, RNAs are biodegradable and short-lived, thus allowing temporary expression and circumventing genome integration^25,27^. Compared to viral vectors, lipid-mediated delivery is superior in terms of larger payload capacity and ease of preparation in addition to its lower immunogenicity and toxicity and higher safety profiles^28,29^. Extensive literature has documented the therapeutic potential of LNP-mediated delivery platforms in immunotherapy^30–32^. After screening a library of ionizable LNPs, we identified 306O_10_ as a promising candidate based on its previously demonstrated efficacy in mRNA-based immunotherapy and CRISPR/Cas9 delivery^33–37^. 306O_10_ LNPs demonstrated lower immunogenicity compared to other formulations and exhibited tropism towards immune cell populations in the spleen and liver, key sites for modulating systemic immune responses^38^.

Initial *in vitro* studies in RAW 264.7 macrophages confirmed efficient 306O_10_-mediated mRNA delivery, evidenced by robust mCherry expression 24 hours post-transfection (Supplementary Fig. 2A). To assess *in vivo* performance and biodistribution, we administered GFP mRNA-loaded particles to C57BL/6 mice via retro-orbital (RO) or tail vein (TV) injection (Supplementary Fig. 2B). Four hours following LNP delivery, we collected lung, liver, and spleen and assessed the transcript levels of GFP. Quantitative RT-PCR analysis revealed widespread GFP mRNA distribution across lung, liver, and spleen tissues, with the RO route yielding the highest transfection efficiency (Supplementary Fig. 2C).

Previous studies have characterized the tropism of 306O_10_ in placental tissue, demonstrating broad distribution across diverse immune and non-immune cell populations^39^. To elucidate the cell type-specific biodistribution of 306O_10_ LNPs, we investigated the cellular uptake patterns within liver and spleen.

To assess cellular targeting in the liver, we evaluated GFP-306O_10_ uptake in primary cultures of liver sinusoidal endothelial cells (LSECs), Kupffer cells, and hepatocytes. Quantitative RT-PCR analysis of GFP transcript levels at 48 hours post-transfection revealed substantial uptake in LSECs, Kupffer cells, and hepatocytes (Supplementary Fig. 2D).

To characterize the immunological targeting profile of 306O_10_ in spleen, we employed the experimental design outlined in Supplementary Fig. 2E. GFP mRNA-encapsulated LNPs were administered via intravenous injection to C57BL/6J mice. Spleen tissues were harvested 4 hours post-administration for subsequent immune cell isolation and quantitative analyses. Magnetic-activated cell sorting (MACS) was employed to isolate CD19^+^, CD11c^+^ and CD11c^-^ populations from the harvested tissues. The level of LNP uptake within each sorted immune subset was then quantified by qRT-PCR measurement of GFP mRNA (Supplementary Fig. 2F). These findings highlight the versatility of 306O_10_ LNPs in targeting diverse immune cell populations, a critical feature for immunomodulatory applications.

### ZF-based repression of the *Myd88* locus achieves efficient immunomodulation under both homeostatic and LPS-induced inflammatory conditions

Having validated our ZF-based repressor *in vitro* and optimized its delivery via 306O_10_ LNPs, we set out to determine whether systemic administration of ZFR11-306O_10_ could effectively repress endogenous *Myd88* expression in wild-type C57BL/6 mice. To test this, an experiment was devised as illustrated in Supplementary Fig. 3A. Mice received a single intravenous injection of either ZFR11-mCherry-306O_10_ or control groups (receiving mCherry-306O_10_ or no LNP/PBS). Six hours post-administration, euthanasia was performed, and blood, lung, and spleen were harvested for RT-qPCR measurement of *Myd88* mRNA levels. We observed *Myd88* repression in ZFR11-mCherry-306O_10_ treatment group across multiple tissues, with transcript levels reduced by 50% in the spleen, 46% in blood, and 16% in lung compared to mCherry-306O_10_ controls (Supplementary Fig. 3B). Additionally, *Myd88* repression was accompanied by a concomitant downregulation of key downstream inflammatory mediators, such as *Icam-1*, *Tnf-α*, *Ncf*, *Il6*, *Ifn-α*, *Ifn-β*, *Ifn-γ, Il-1β, and Stat4* (Supplementary Fig. 3C).

We next investigated whether LNP-mediated delivery of the *Myd88* repressor could modulate inflammatory responses in a lipopolysaccharide (LPS)-induced septicemia model, a clinically relevant system manifesting elevated *Myd88* expression^40–42^. This acute inflammatory condition is particularly relevant for evaluating *Myd88*-targeted interventions, as septicemia remains a significant global health burden with limited therapeutic options due to its complex pathophysiology involving dysregulated inflammatory cascade^43,44^

C57BL/6 mice received intraperitoneal injection of LPS followed by intravenous administration of ZFR11-306O_10_ two hours later (Fig. 1A). Remarkably, 24 hours post-LNP treatment, we observed robust *Myd88* repression in blood (79%), lung (58%), and liver (22%) compared to PBS-treated controls (Fig. 1B). This repression effectively prevented the LPS-induced upregulation of multiple inflammatory mediators downstream of *Myd88* signaling, including *Icam-1*, *Tnf-α*, *Ncf*, *Il6*, *Ifn-α*, *Ifn-β*, *Ifn-γ*, and *Stat4* (Fig. 1C-E). Analysis of plasma cytokine concentrations by quantitative chemiluminescent ELISA revealed a trend towards lower level of cytokines in *Myd88*-repressed mice and further corroborated the anti-inflammatory effects of ZF-based *Myd88* repression (Fig. 1F). Collectively, these results demonstrate that LNP-mediated delivery of ZFR11 can effectively modulate *Myd88* expression under both homeostatic and inflammatory conditions, establishing proof-of-concept for this synthetic biology approach to immunomodulation.

**Figure 1.**
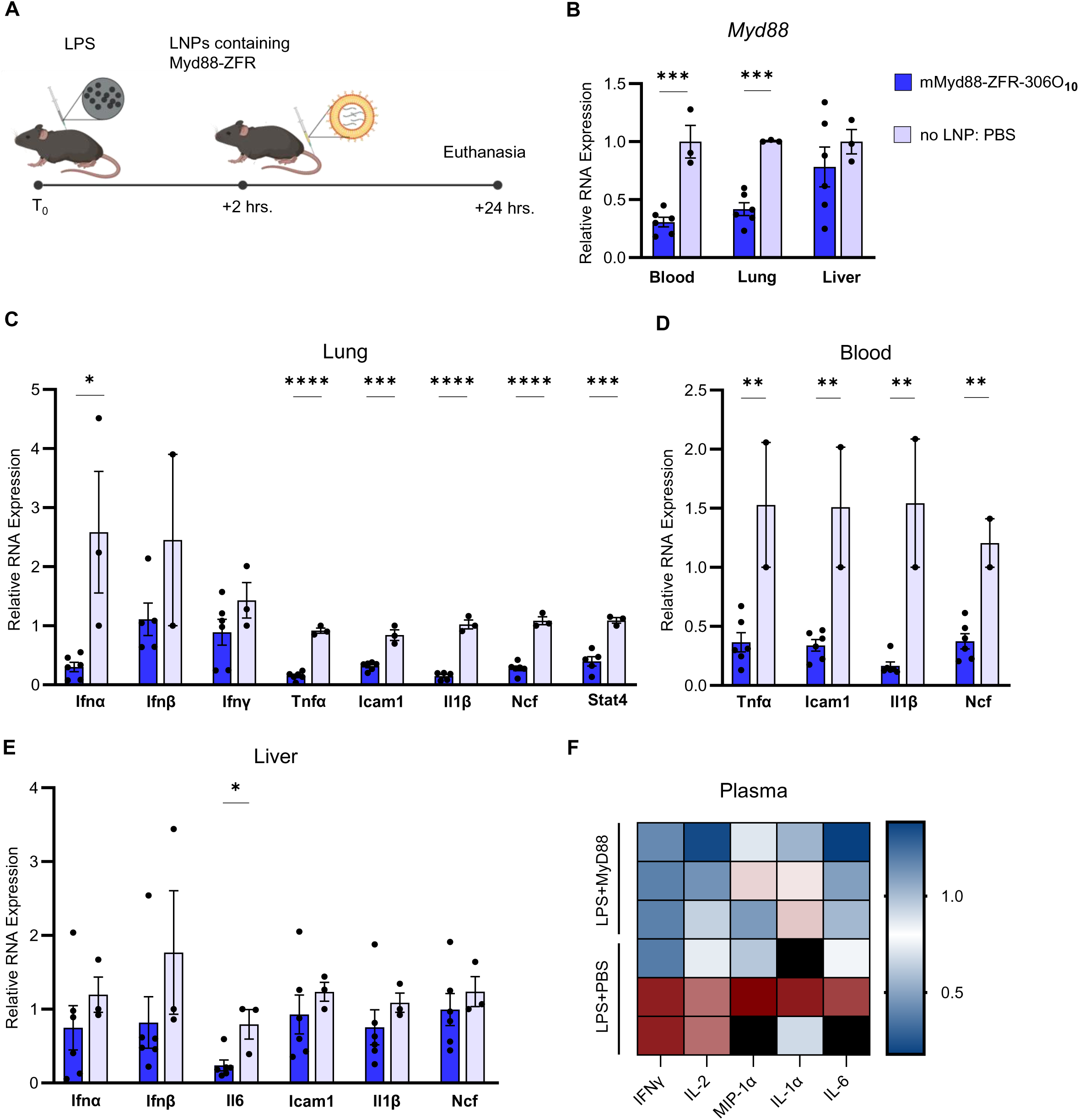
Delivery of lipid nanoparticles carrying mRNA encoding *Myd88*-targeting zinc finger enables immunomodulation during LPS-induced inflammation. (A) Schematic representation of experimental design. C57BL/6 mice received intraperitoneal injection of LPS (2.5 mg kg^-1^) followed by tail vein administration of either ZFR11-306O_10_ or no LNP/PBS (control) 2 hours later. Tissues were collected 24 hours post-LNP administration. (B) qRT-PCR analysis of *Myd88* expression in blood, lung, and liver tissues following LPS challenge and zinc finger-mediated repression (N = 6 mice for ZFR11-306O_10_ group, N = 3 for PBS control group). Data are presented as mean + s.e.m. (C) qRT-PCR analysis of inflammatory gene expression (*Ifn-α*, *Ifn-β*, *Ifn-γ*, *Icam-1*, *Ncf*, *Tnf-a*, *Il-1β*, and *Stat4*) in lung. Expression levels were normalized to PBS-treated control mice (N = 6 for ZFR11-306O_10_ group, N = 3 for PBS group). Data are presented as mean + s.e.m. (D) qRT-PCR analysis of inflammatory gene expression (*Tnf-a*, *Icam-1, Il-1β*, and *Ncf*) in blood. Expression levels were normalized to PBS-treated control mice (N = 6 for ZFR11-306O_10_ group, N = 3 for PBS group). Data are presented as mean + s.e.m. (E) qRT-PCR analysis of inflammatory gene expression (*Ifn-α*, *Ifn-β*, *Il6*, *Icam-1*, *Il-1β*, and *Ncf*) in liver. Expression levels were normalized to PBS-treated control mice (N = 6 for ZFR11-306O_10_ group, N = 3 for PBS group). Data are presented as mean + s.e.m. (F) Measurement of a panel of inflammatory cytokines in plasma using a multiplex-ELISA assay; values are displayed in the heatmap as relative measured concentration (pg/mL) compared to PBS-treated control mice. : IFNγ (N = 2 per group), IL2 (N = 3 for ZFR11-306O10, N = 2 for PBS), MIP-1a (N = 3 for ZFR11-306O_10_, N = 2 for PBS), IL-1a (N = 3 for ZFR11-306O_10_, N = 2 for PBS), and IL-6 (N = 3 for ZFR11-306O_10_, N = 2 for PBS). IFNγ, interferon gamma; IL, interleukin; MIP, macrophage inflammatory protein. Statistical analysis was performed using the unpaired parametric t-test. * *P* ≤ 0.05 was considered statistically significant.

### ZFR11 functions as an anti-adjuvant to modulate AAV-directed humoral immunity and enhance the efficiency of viral-based gene delivery

To mount an effective adaptive immune response, antigen-presenting cells must activate T cells through both antigen presentation and co-stimulatory signals at the immunological synapse. This principle is leveraged in vaccines through the use of adjuvants, which enhance immune activation. A critical component of these stimulating signals includes inflammatory cytokines, whose production is regulated by NF-κB signaling, with Myd88 serving as a key upstream mediator. We predicted that selective *Myd88* repression during antigen exposure could function as an ‘anti-adjuvant,’ attenuating inflammation at the immunological synapse and preventing immune activation. We therefore investigated whether our *Myd88* repressor could regulate humoral adaptive immunity against AAV vectors in pre-immunized hosts, addressing a major challenge in AAV-based gene therapy re-dosing. This anti-adjuvant approach has shown promise in various therapeutic applications, including allergy immunotherapy, transplant tolerance, and autoimmune disease treatment^45–48^.

To investigate the effect of *Myd88* repression in hosts with pre-existing anti-AAV antibodies, we established an AAV2-pre-exposed mouse model (Figure 2A) using AAV2, an FDA-approved gene therapy vector. C57BL/6J mice received two retro-orbital (RO) injections of AAV2-GFP to induce pre-existing immunity. We developed a reverse vaccination strategy using LNP-306O_10_-encapsulated mRNA encoding three key components: ZFR11 *Myd88* suppressor (anti-adjuvant), VP1 (AAV2 capsid antigen), and mCherry (gene therapy cargo). Mice were divided into three groups (N = 10 per group) receiving four weekly doses of either: ZFR11-VP1-mCherry (group 1), VP1 alone (group 2), or PBS control (group 3). Two weeks after the final vaccination, all mice received a single intravenous co-injection of AAV2-mCherry and ZFR11-306O_10_. Analysis was performed three weeks post-AAV2-mCherry challenge, followed by collection of blood and tissue samples for downstream processing.

**Figure 2.**
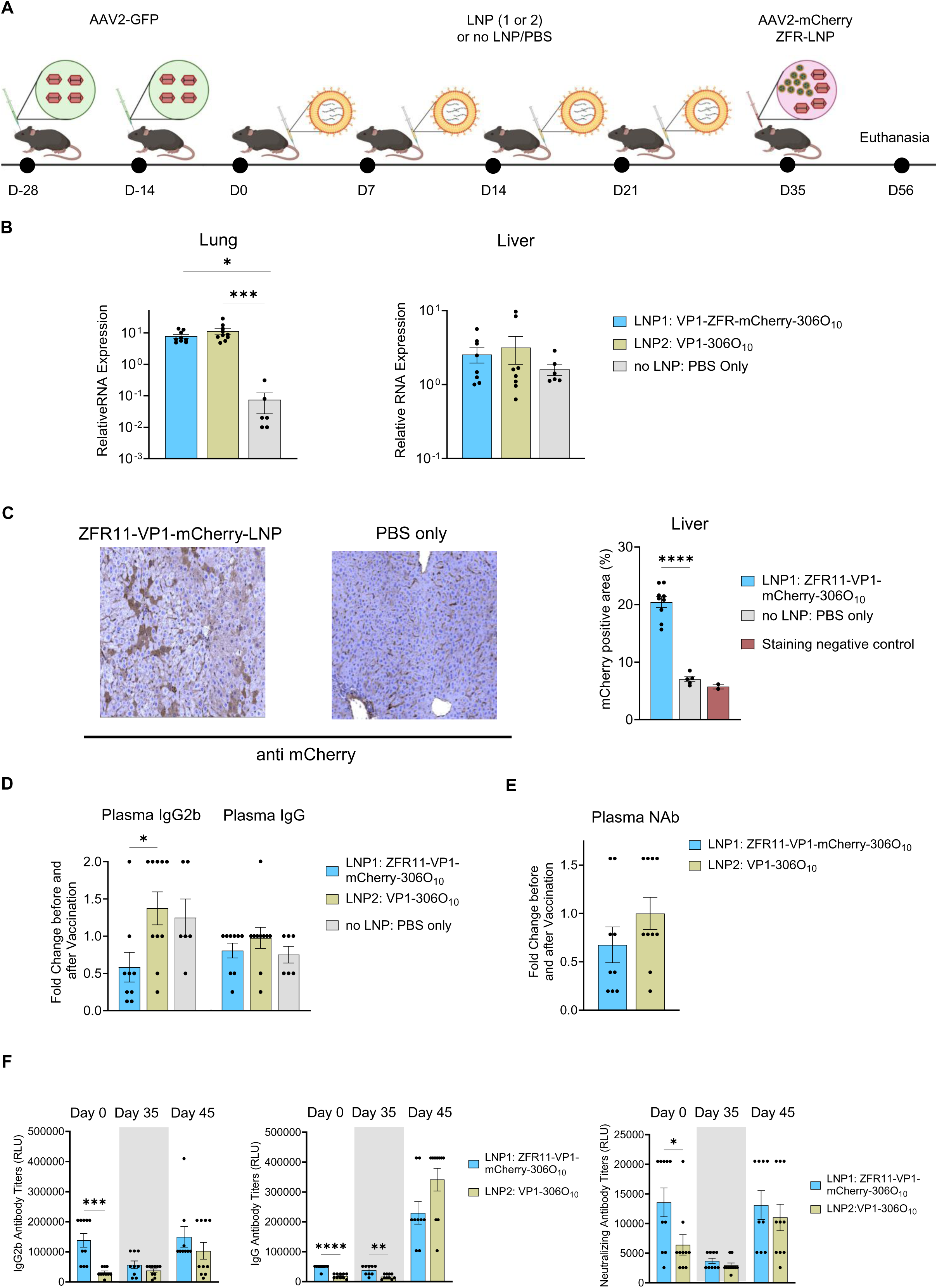
Co-delivery of *Myd88* repressor ZFR11 and gene therapy related antigen modulates humoral immunity in AAV pre-immunized mice. (A) Schematic representation of experimental design. C57BL/6J mice received two retro-orbital injections of AAV2-GFP (5 × 10^11^ GC) two weeks apart. Two weeks later, mice were administered weekly doses of LNP-encapsulated mRNA (0.7 mg kg-1) encoding either ZFR11-VP1-mCherry (N = 10), VP1 only (N = 10), or no LNP/PBS control (N = 6) for four weeks. Two weeks after the final vaccination, mice received a single intravenous co-injection of AAV2-mCherry (5 × 10^11^ GC) and ZFR11-LNP (0.7 mg kg-1). Analyses were performed three weeks post-AAV2 challenge. (B) qRT-PCR analysis of *mCherry* expression in liver and lung tissues (N = 8-10 mice per treatment group, N = 6 for PBS). Data are presented as mean + s.e.m. Statistical analysis was performed using one-way ANOVA followed by Dunnett’s multiple comparisons test. \**P* ≤ 0.05 was considered statistically significant (C) Left: Representative immunohistochemical staining for mCherry in liver sections. Right: Quantification of mCherry staining intensity (ZFR11-VP1-mCherry-306O_10_: N = 9, PBS: N = 5, staining negative control/mCherry uninjected: N = 2). Data are presented as mean + s.e.m. Statistical analysis was performed using one-way ANOVA followed by Dunnett’s multiple comparisons test. \**P* ≤ 0.05 was considered statistically significant. (D) ELISA measurement of plasma anti-AAV2 total IgG and IgG2b levels. Results shown as fold change in optical density at day 35 (two weeks post-vaccination) relative to day 0 (pre-vaccination baseline) (N = 9-10 per treatment group, N = 6 for PBS). Statistical analysis was performed using one-way ANOVA followed by Dunnett’s multiple comparisons test. \**P* ≤ 0.05 was considered significant. (E) ELISA measurement of serum anti-AAV2 neutralizing antibody levels showing a 32% reduction in the ZFR11-VP1-mCherry group compared to VP1 control at day 35 (two weeks post-vaccination) relative to day 0 (pre-vaccination baseline) (N = 9-10 mice per treatment group, N = 6 mice for PBS). Statistical analysis was performed using the unpaired parametric t-test. \**P* ≤ 0.05 was considered significant. (F) Time-point analysis of plasma total IgG, IgG2b, and neutralizing antibodies titers in Relative Light Units (RLU) at day 0 (pre-vaccination baseline), day 35 (two weeks post-vaccination), and day 45 (10 days after AAV2-mCherry rechallenge) (N = 9-10 per treatment group).

We first assessed whether vaccination with *Myd88* repressor could influence the efficiency of AAV-mediated transgene delivery mimicking clinical scenarios requiring therapeutic gene delivery in AAV pre-exposed populations. Quantitative RT-qPCR analysis identified significantly elevated mCherry (AAV cargo transgene) transcript levels in lung tissues of the mice receiving ZFR11-VP1-mCherry and VP1 vaccinations compared to control group receiving PBS alone (∼100-fold and ∼150-fold, respectively) (Fig. 2B). A similar trend was observed in the liver, albeit with more modest increases (∼1.6-fold and ∼2-fold, respectively). In agreement with earlier qRT-PCR results, quantification of the non-fluorescent IHC images confirmed a ∼3-fold higher mCherry protein expression in liver sections from ZFR11-VP1-mCherry-vaccinated mice versus PBS control (Fig. 2C).

Next, we investigated whether treatment with ZFR11-VP1-mCherry-306O_10_ LNPs could attenuate AAV-specific humoral responses by measuring both immunoglobulin G (IgG) levels and neutralizing antibody titers. To assess humoral immunity, we quantified anti-AAV2 IgG and IgG2b levels in plasma samples by enzyme-linked immunosorbent assay (ELISA). We observed that ZFR11-VP1-mCherry treatment significantly reduced anti-AAV2 IgG2b levels by 58% compared to VP1 controls before (day 0) and after vaccination (day 35) (Fig. 2D). This group also showed a trend toward lower IgG responses, with levels 18% below VP1 controls (Fig. 2D). Additionally, neutralizing antibody levels were reduced by 32% in ZFR11-VP1-mCherry vaccinated animals compared to VP1 controls before (day 0) and after vaccination (day 35) (Fig. 2E). While vaccination with ZFR11-VP1-mCherry initially reduced IgG2b and neutralizing antibody levels (comparing days 28 and 63, before and after vaccination), this tolerance effect was transient. AAV2-mCherry rechallenge at two weeks post-injection (day 73) restored these levels to those observed after initial AAV2-GFP administration (Fig. 2F).

Notably, the therapeutic intervention prevented further upregulation beyond baseline, representing this strategy’s efficacy in maintaining stable antibody responses [44]. These results underscore the immunomodulatory effects of ZFR11-VP1-mCherry as an anti-inflammatory agent, suggesting its capacity to modify established immune responses even in the context of pre-existing immunity, though further optimization of dose, timing, and frequency is needed (Figure 2F).

## Discussion

Our work establishes a novel epigenetic editor for targeted immunomodulation using zinc finger-based transcriptional repressors delivered via lipid nanoparticles. Exploiting this modality, we demonstrated efficient repression of endogenous Myd88 both *in vitro* and *in vivo*, effectively dampening inflammatory responses during LPS-induced model of septicemia and enhancing outcomes in AAV gene therapy applications (Fig. 3). This strategy addresses key limitations of current immunomodulatory approaches by providing precise transient control over immune responses.

**Figure 3.**
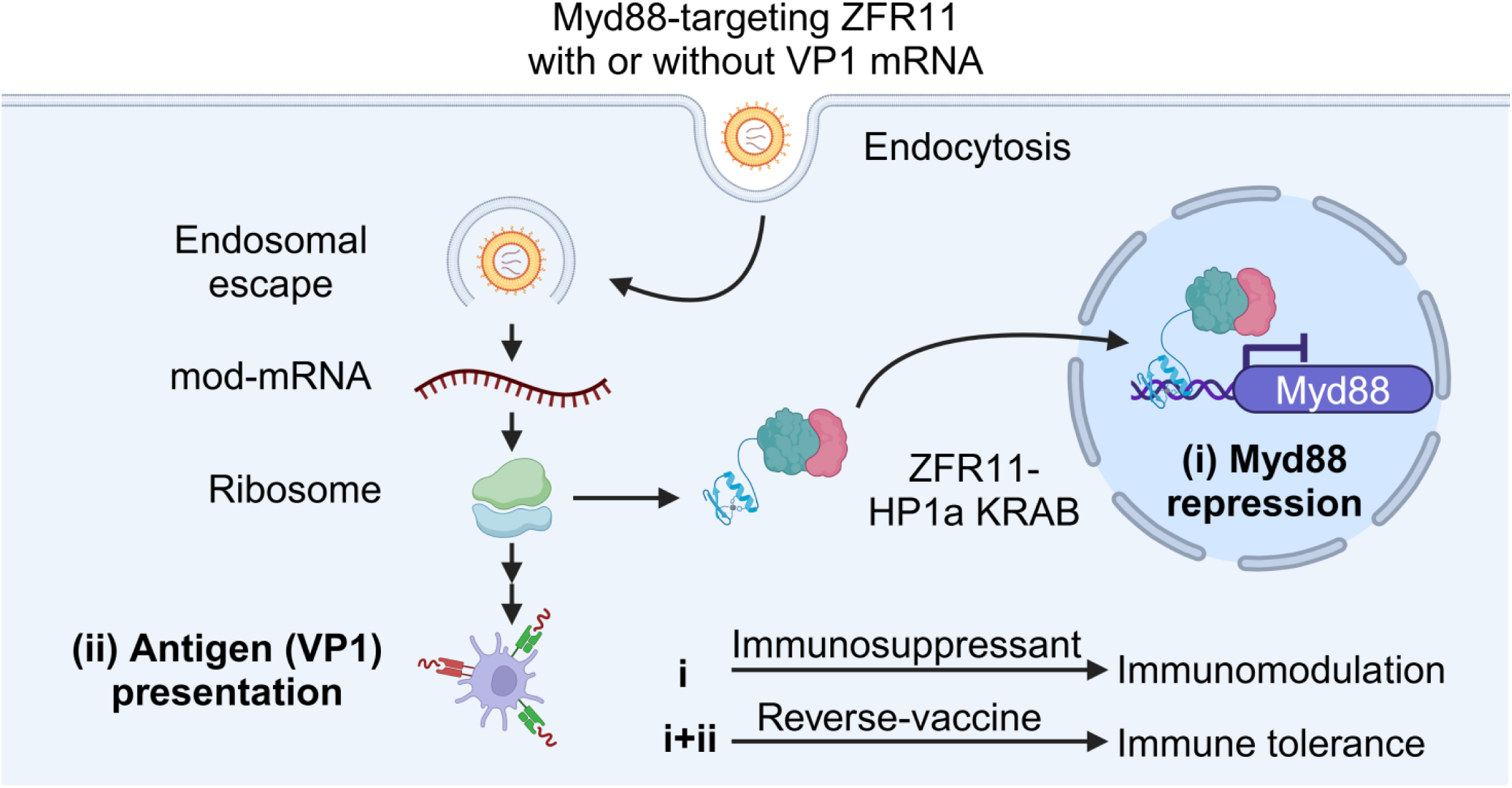
Schematic representation of the dual therapeutic applications of Myd88-targeting ZFR11. The lipid nanoparticle-mediated delivery of ZFR11-HP1a-KRAB mRNA enables two distinct therapeutic approaches: (i) Myd88 repression alone acts as an anti-inflammatory agent, a more targeted immunosuppressant, to modulate systemic inflammation in conditions such as septicemia, and (i+ii) co-delivery with antigen-encoding mRNA (e.g., AAV2 VP1) in an immunosilent formulation serves as a reverse-vaccine strategy to induce antigen-specific immune tolerance. Following endocytosis and endosomal escape, the modified mRNA is translated to produce either ZFR11 alone or in combination with the target antigen (VP1). ZFR11-HP1a-KRAB localizes to the nucleus where it binds the Myd88 promoter region to facilitate epigenetic repression. When delivered alone (i), this leads to targeted immunomodulation through reduced inflammatory signaling. When co-delivered with antigen (i+ii), the combination of Myd88 repression and antigen presentation promotes the development of antigen-specific tolerance, as demonstrated by enhanced AAV2 vector readministration capacity.

Our approach, which uses an effective LNP delivery system to deliver zinc finger repressor-encoding mRNAs, represents a significant advance over existing methods. Unlike viral vectors or small molecule inhibitors, our work combines the specificity of zinc finger proteins with the versatility of 306O_10_ lipid nanoparticles to enable transient targeting of immune cell populations for short-term immunomodulation ^38,39^. The scalability and relatively straightforward manufacturing process of our LNP-based delivery system provides advantages for clinical translation. Whereas viral vectors often face production challenges, LNP-based therapeutics can be manufactured using well-established processes.

Our data demonstrate two distinct therapeutic applications of this platform. In the context of acute inflammation, ZFR11-mediated *Myd88* repression effectively prevents the cascade of inflammatory signaling typically associated with septicemia. This suggests potential applications in treating various inflammatory conditions where precise, temporary immune suppression is desired.

In the context of AAV gene therapy, our findings usher in a new era for mitigating pre-existing immunity. The ability to temporarily suppress anti-AAV antibody production through *Myd88* repression provides a viable solution to one of the field’s most significant challenges. While the effect is transient, the window of reduced immunity could be sufficient to enable successful vector readministration.

Future therapeutic applications of this platform may benefit from several potential optimizations. Alternative dosing schedules, modified LNP formulations, or combinations with existing immunomodulatory approaches could extend the duration of effect. Additionally, the modular nature of our system allows for targeting of other immune regulators beyond Myd88, potentially enabling treatment of diverse immunological conditions. Further investigations can unveil the mechanisms underlying the observed immunomodulatory effects and explore applications across different disease models to determine the broader utility of this epigenetic engineering platform.

In conclusion, our findings highlight the potential of combining precisely tuned gene control modalities with enhanced mRNA delivery strategies to address diseases characterized by dysregulated inflammatory pathways. Our study opens new possibilities for achieving precise, safe, and effective modulation of immune responses across a broad spectrum of diseases.

## Methods

### Zinc Finger *Myd88* repressor fusion DNA constructs

A panel of 16 zinc finger (ZF)-based transcriptional repressors targeting the *Myd88* promoter was designed and constructed through a multi-step cloning process. First, the repressive domains (HP1a and KRAB) were PCR-amplified from previously constructed vectors described in our earlier work^22^.These domains were then fused via overlap extension PCR to generate the HP1a-KRAB DNA fragment. The resulting fragment was inserted into an L1L2 entry backbone using BsaI-based Golden Gate cloning (NEBridge Golden Gate Assembly Kit, Cat. No. E1601S). A library of 16 distinct zinc finger DNA fragments, each targeting different regions of the *Myd88* promoter, was then cloned upstream of the HP1a-KRAB using In-Fusion cloning technology (Takara Bio’s In-Fusion Snap Assembly Master Mix, Cat. No. 638948).The final expression constructs were assembled through a three-fragment Gateway recombination reaction (ThermoFisher Scientific, Cat. No. 12538120) utilizing L4R1-CAG (promoter), L1L2-ZF-HP1a-KRAB (repressor), and R1R2-LVGTW3 (terminator) constructs. This approach generated 16 distinct ZF repressor DNA constructs. Complete sequences of all zinc finger arrays and the ZFR1-HP1a-Krab fusion DNA fragment are provided in Supplementary Table 1.

### Optimization of ZF-based *Myd88* Repressor for enhanced efficacy and reduced immunogenicity

To maximize the therapeutic potential of our ZF-based *Myd88* repressor while minimizing undesired immune responses, we implemented a series of strategic design optimizations. These modifications encompassed both the mRNA component and the choice of epigenetic modulator. For the mRNA component, we incorporated N1-methyl pseudouridine nucleoside modifications and employed a uridine depletion strategy. N1-methyl pseudouridine has been shown to enhance mRNA stability and reduce recognition by pattern recognition receptors, thereby attenuating innate immune activation^49–52^. Uridine depletion further contributes to reduced immunogenicity by minimizing uridine-rich sequences that can trigger Toll-like receptor activation^53^.

We also optimized the nucleic acid sequence for mammalian codon usage, a strategy that has been demonstrated to improve translational efficiency and potentially reduce immunogenicity by mimicking endogenous mRNA characteristics^54,55^. To eliminate double-stranded RNA (dsRNA) contaminants, which are potent activators of innate immune responses, we employed a cellulose-based dsRNA removal process in our mRNA preparation protocol^56,57^.

The selection of zinc finger proteins as our epigenetic modulator was informed by their favorable immunological profile and structural characteristics. Compared to other gene-editing technologies such as CRISPR-Cas9, zinc finger proteins exhibit lower immunogenicity and have a more compact structure, potentially facilitating delivery and reducing the likelihood of eliciting an immune response^58^.

These multifaceted optimizations were designed to synergistically enhance the efficacy of our ZF-based *Myd88* repressor while minimizing its potential to trigger undesired immune responses, thereby improving its therapeutic index and translational potential.

### mRNA and lipids

The following mRNAs and reagents were used for nanoparticle formulation: CleanCap® modified mRNAs (encoding GFP, mCherry, ZFR11, and VP1) from TriLink Biotechnologies; Cholesterol from Sigma-Aldrich (catalog number C8667, ≥99% pure, extracted from sheep wool); and phospholipids from Avanti Polar Lipids including DOPS (sodium 1,2-dioleoyl-sn-glycero-3-phospho-L-serine, catalog number 840035P) and C14-PEG2000 (ammonium 1,2-dimyristoyl-sn-glycero-3-phosphoethanolamine-N-[methoxy(polyethylene glycol)-2000], catalog number 880150P).

### Lipid Nanoparticle Formulation

Lipid nanoparticles were prepared through a modification of the ethanol injection technique. The initial step involved dissolving the component lipids (lipidoids, helper lipids, cholesterol, and C14-PEG2000) in ethanol, with each component at 1-10 mg/mL concentration. These lipid components were combined at specific molar proportions: lipidoids (35), helper lipids (40), cholesterol (22.5), and PEG-lipid (2.5). This lipid solution was then combined with mRNA that had been prepared in 10 mM sodium citrate buffer (pH 3.0), maintaining a lipidoid to mRNA weight ratio of 10:1. Following particle formation, residual ethanol was removed by dialyzing the LNP suspension against PBS using Thermo Scientific cassettes with a 3 kDa molecular weight cutoff.

### AAV vectors

Cas9 plasmid was purchased from Addgene (AAV-CMVc-Cas9 #106431). mCherry construct was a premade AAV vector purchased from PackGene Biotech, (ssAAV-CAG-mCherry.WPRE.SV40pA, Cat. No. AAV-EA022).

### AAV packaging and purification

AAV plasmid integrity was verified through SmaI restriction enzyme digestion, specifically examining ITR regions. Following validation, these constructs served as templates for AAV2-Cas9 and AAV2-mCherry viral production by PackGene Biotech, LLC. Viral titers were determined using Real-time SYBR Green PCR against standard curves prepared from linearized parental AAV vectors, establishing concentrations of 1.5 × 10^13^ GC/ml.

### Cell culture

Neuro-2a cells (purchased from ATCC) were cultured at 37°C in a 5% CO_2_ environment. The culture medium consisted of Dulbecco’s modified Eagle’s medium (DMEM) purchased from Life Technologies supplemented with the following components: FBS (10%, Life Technologies), sodium pyruvate (1.0 mM, Life Technologies), glutamine (2 mM), and a streptomycin-penicillin antibiotic mixture (1%, Gibco).

### Transfection of *in vitro* cultured cells

For *in vitro* transfection studies, Neuro-2a cells we seeded in 24-well plates at a density of approximately 50,000 cells per well. The following day, transfection was using Lipofectamine LTX to deliver multiple plasmid components: Cas9 nuclease (50 ng), gRNA (10-100 ng), ZFR (150 ng), dCas9-HP1a-KRAB (100 ng), YFP for monitoring transfection efficiency (25 ng), and a puromycin resistance marker (50 ng). 24 hours following transfection, puromycin selection was done using 0.5 μg/ml concentration (Gibco-life tech)

### Quantitative RT-PCR (qRT-PCR) analysis

For RNA extraction, cell lysis was performed, and RNA was extracted using either Life Technologies’ Trizol or Qiagen’s RNAEasy Plus Mini Kit. The resulting RNA underwent reverse transcription to cDNA using Thermo Fisher’s High-Capacity RNA-to-cDNA Kit. Quantitative PCR analysis employed SYBR Green PCR Master Mix (Thermo Fisher), with 18S rRNA serving as the normalization standard. We calculated relative expression changes using the 2^−ΔΔCt^ method, comparing against control group values. The complete list of primers used for quantitative PCR are in Supplementary Table 2.

### Plasma biomarker analysis of liver function

The isolation of plasma from blood samples was achieved by centrifugation, performed for 10 minutes at 2,000 × g while maintaining 4°C temperature. Plasma bile acids and alanine aminotransferase (ALT) levels were analyzed by IDEXX Laboratories (IDEXX Reference Laboratories, USA). Analysis was performed using the IDEXX Catalyst chemistry analyzer system. Total bile acids were measured through an enzymatic cycling reaction utilizing 3α-hydroxysteroid dehydrogenase (reported in µmol/L). ALT activity was determined via a coupled enzymatic assay monitoring the rate of NADH oxidation spectrophotometrically (reported in IU/L). All assays were performed with appropriate calibration controls following manufacturer’s validated protocols.

### ELISA-based chemiluminescent assay for plasma cytokine analysis

Multiplexed analysis of plasma cytokines was conducted at the UPMC Cancer Proteomics Facility: Luminex Core Laboratory using two magnetic bead-based immunoassay panels. A Mouse High Sensitivity T Cell 8-plex kit (MHSTCMAG-70K-08, Millipore) was used to measure IL-6, IL-4, IL-2, IL-1β, IL-1α, IL-17A, IL-10, and IFN-γ from 60 μL plasma samples. MIP-1α levels were determined using the Mouse Cytokine MAGNETIC Panel 1 (MCYTOMAG-70K-01) from 35 μL plasma samples. All samples were analyzed in duplicate following manufacturer’s protocols. Briefly, samples and calibrators were incubated with analyte-specific antibody-conjugated magnetic beads in 96-well plates. After washing, biotinylated detection antibodies were added followed by streptavidin-horseradish peroxidase. Bead fluorescence was measured using a Luminex analyzer and cytokine concentrations were calculated against standard curves using xPONENT software.

### IFN-γ ELISPOT Assay

IFN-γ ELISPOT assays were performed by the Immunology Core at the Gene Therapy Program, Perelman School of Medicine, University of Pennsylvania. Briefly, AAV2 capsid peptide library (Mimotopes, Victoria, Australia) consisting of 145 peptides was synthesized as 15-mers with 10 amino acid overlap. The library was subdivided into three peptide pools (A, B, and C) at approximately 2 mg/mL per peptide. Cryopreserved mouse splenocytes were analyzed for T-cell responses to AAV2 capsid using murine IFN-γ ELISPOT. Assay wells were stimulated with DMSO (negative control), individual AAV2 capsid peptide pools, and PMA/ION (positive control). For individual peptide pools and DMSO control, 2.0 × 105 cells were plated per well, with data calculated and presented as spot forming units (SFU) per million cells. For the PMA/ION positive control, 2.0 × 103 cells were plated per well. All samples were run in duplicate. A response was considered positive if it met both criteria: average value ≥ 26 SFU/million cells and at least three times greater than the DMSO negative control value. ELISPOT data were analyzed using a CTL Immunospot S6 Core Analyzer.

### Anti-AAV2 Antibody Assays

Anti-AAV2 antibodies were quantified utilizing both *in vitro* neutralization and ELISA assays at the Gene Therapy Program’s Immunology Core facility at the University of Pennsylvania’s Perelman School of Medicine (Philadelphia, PA).

### IgG and IgG2B ELISA

The immunoassay utilized Nunc maxisorp plates (Thermo Fisher Scientific, Waltham, MA) with an initial AAV2 particle coating step (2 × 10^10^ particles per mL in carbonate buffer, pH 9.6) performed overnight at 4°C. Standard curves were generated using Sigma-Aldrich (St. Louis, MO) purified immunoglobulins (IgG and IgG2B) in serial dilutions. Following a room temperature blocking step (1 hour) with PBS containing 2% BSA and 0.05% Tween-20, we applied diluted plasma samples in duplicate wells for a 3-hour room temperature incubation. Detection employed two horseradish peroxidase (HRP)-conjugated secondary antibodies from Southern Biotech: anti-mouse IgG-HRP (1:20,000) and anti-mouse IgG2B-HRP (1:10,000). After incubating at 37°C for 1 hour and washing, we developed the plates using SIGMAFASTTM OPD substrate (Sigma-Aldrich) according to manufacturer specifications, with absorbance readings taken at 492 nm.

### Neutralizing Antibody Assay

Anti-AAV2 neutralizing antibody levels were evaluated in selected plasma specimens using a cell-based *in vitro* approach. The protocol starts with seeding HEK293 cells (1×10^5^ cells/well) into 96-well plates, allowing 24 hours for attachment. Test samples were then prepared by heat inactivation followed by serial dilution, combining them with luciferase-expressing AAV2 vector (1×10^4^ viral particles/cell) for a 1-hour incubation at 37°C. This mixture was introduced to the cells for a 24-hour incubation period. Using the Galacto-Star System (Applied Biosystems), luciferase activity was measured. The neutralizing antibody titer was defined as the maximum sample dilution that achieved at least 50% reduction in luciferase expression compared to controls without inhibition. For reference, a 1:10 neutralizing antibody titer indicates that a sample diluted ten-fold exhibits a luciferase signal below 50% of the non-inhibition control value.

### Animals

Our animal research protocols adhered to established laboratory animal care and usage guidelines, with full approval from the University of Pittsburgh’s Institutional Animal Care and Use Committee (IACUC). All experiments followed institutional protocols and included both male and female C57BL/6 mice (purchased from JAX, Stock #000664) aged 6-8 weeks. Each experimental group contained a minimum of three animals, with precise group sizes documented in the corresponding figure legends. C57BL/6 mouse strain was utilized for both AAV LPS and experiments.

### Retro-orbital injections

For administration of AAV particles, the retro-orbital route targeting the venous sinus was selected. Mice were anesthetized using isoflurane at 3% concentration. 100 microliters of AAV solution, containing between 1 × 10^11^ and 1 × 10^12^ genome copies per animal, was injected into each mouse’s left eye.

### Tissue harvest

Using CO_2_ inhalation for euthanasia, tissue samples were collected from multiple organs - liver, spleen, lung, and bone marrow - as well as blood samples. Each tissue sample was immediately placed in RLT Plus buffer from Qiagen, followed by preservation through snap freezing methods for later RNA extraction and analysis.

### *In vivo* LPS Administration

*Escherichia coli* strain 0127: B8 derived lipopolysaccharides were administered via intraperitoneal (i.p.) route (LPS purchased from Sigma-Aldrich, St. Louis, MO, USA). The LPS was prepared as a 2.5 mg/ml solution in PBS. At 26 hours following LPS administration (as depicted in the experimental timeline schematics), animals were euthanized using CO2 inhalation.

### Statistical analysis and reproducibility

All *in vitro* experiments were conducted in triplicate, with consistency observed across replicates. For animal studies, a minimum of three biological replicates were performed, yielding reproducible results. The assignment of mice to experimental or control groups was performed randomly, though experimenters were not blinded during data acquisition or analysis. Data are presented as the mean + s.e.m. The variable N denotes either the number of individual transfections (for *in vitro* work) or the number of animals (for *in vivo* studies). Statistical analyses were performed using GraphPad’s Prism 10 Software, which are detailed within figure legends. When comparing two groups for statistical significance, two-tailed unpaired t-tests was employed, while multiple group analyses was performed by one-way ANOVA with Dunnett’s multiple comparisons test. *\*P* ≤ 0.05 was considered significant (with additional thresholds at \*\**P* ≤ 0.01, \*\*\**P* ≤ 0.001, \*\*\*\**P* ≤ 0.0001).

## Data Availability

The raw and processed data supporting the findings of this study are available from the corresponding author subject upon reasonable request. All materials are available upon completion of a material transfer agreement.

## Acknowledgements

This work was supported by sponsored research agreements from the Cystic Fibrosis Foundation through Genexgen Inc. Additional funding was provided by the National Institute of Biomedical Imaging and Bioengineering of the National Institutes of Health under award number R01EB024562, the National Institutes of Health award number DP2-HD098860, and a Startup fund from the Department of Pathology. We gratefully acknowledge Dr. Mo Ebrahimkhani for his valuable guidance and support throughout this project. This study utilized the UPMC Cancer Proteomics Facility: Luminex Core Laboratory, with special thanks to Denise Prosser. We thank Dr. Jessica A. Chichester, Director of the Immunology Core, Gene Therapy Program, Perelman School of Medicine, University of Pennsylvania, for conducting the neutralizing antibody and ELISPOT assays. We also acknowledge Miles Stampo and Amanda Christina Fisher Mihalik at the Division of Laboratory Animal Resources (DLAR) Veterinary Services, University of Pittsburgh, for their support with animal studies. Biorender was used to create several of the schematics shown in this work.

## Author Contributions

T.M. and S.K. designed the study and the associated experiments. T.M., S.K., S.L., and K.A.W. conceived the methodology for experiments and provided strategic guidance. T.M. generated the constructs and performed *in vitro* experiments. T.M., M.N.T., S.A., A.A., V.M., K.K., S.L., and R.L. conducted *in vivo* experiments. T.M. and S.K. analyzed the data. T.M., Y.X., M.N.T., and S.K. performed data visualization and figure preparation. S.K. supervised the study and secured funding. T.M. and S.K. wrote the original manuscript draft. T.M., S.K., M.N.T., and A.K. contributed to manuscript editing and revising the final draft of the manuscript.

---

**Supplementary Figure 1.**
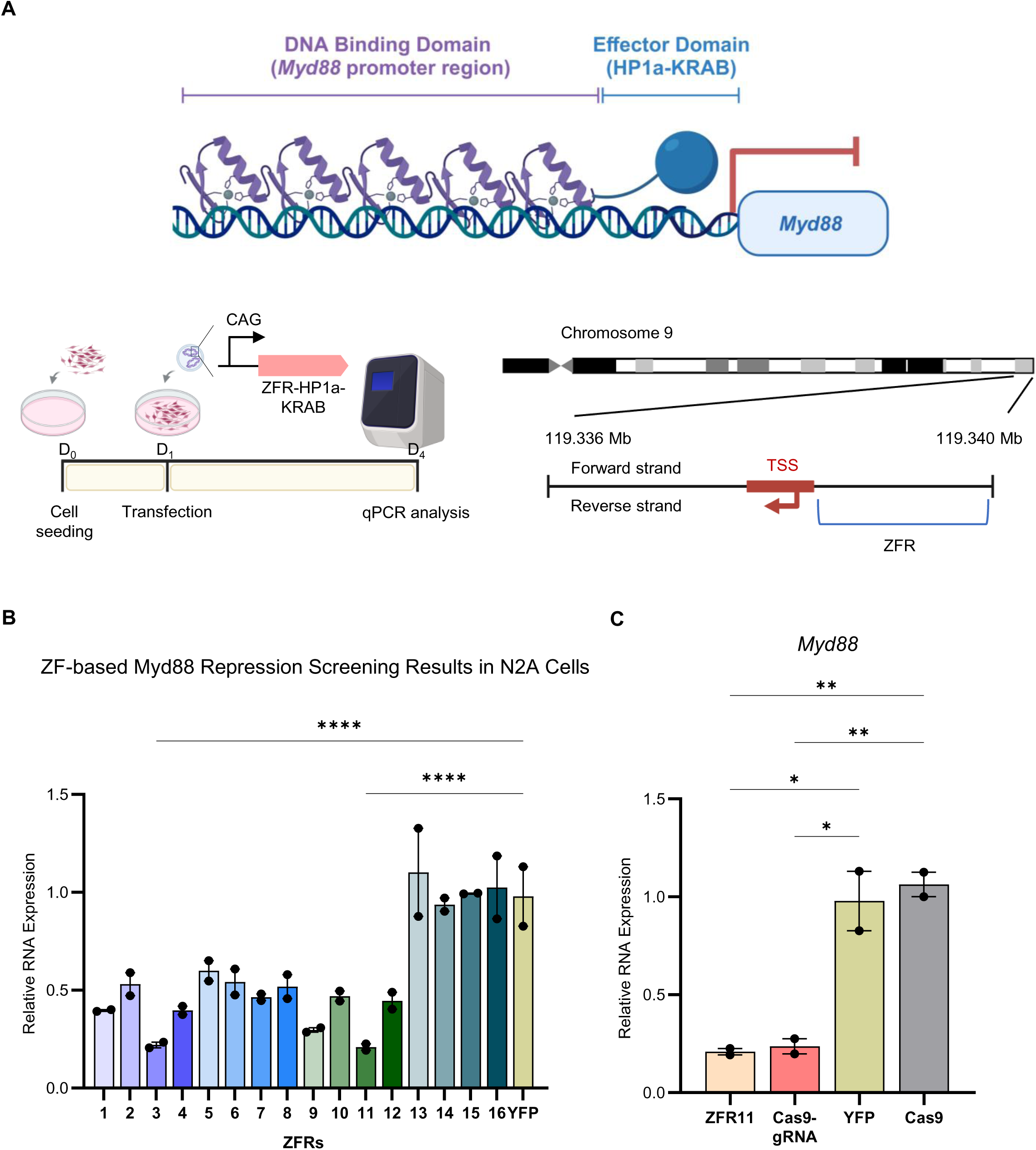
Evaluating zinc-finger HP1a-Krab *Myd88* repression *in vitro.* (A) Top: Schematic of ZF-based artificial transcription factor (ATF) recruitment to the *Myd88* promoter region. ZF-based ATF is composed of an effector domain and a DNA-binding domain. Bottom left: Schematic of the experimental design. Mouse neuroblastoma (N2a) cells were transfected with either one of 16 different *Myd88*-targeting ZF-based repressors or yellow fluorescent protein (YFP)-expressing plasmid as a control. *Myd88* mRNA expression levels were analyzed by qRT-PCR three days post-transfection. Bottom right: Schematic representation of the ZFR11 binding sites within the Myd88 promoter region. (B) Fold changes in Myd88 mRNA transcript levels were quantified relative to the YFP control group (n = 2 biologically independent samples). Data are presented as mean + s.e.m. Statistical analysis was performed using one-way analysis of variance (ANOVA) followed by Dunnett’s multiple comparisons test. Differences were considered statistically significant at \**P* ≤ 0.05. ZFR1 through ZFR12 showed significant differences compared to the YFP control, while ZFR13 through ZFR16 showed no significant differences. For clarity of data presentation, statistical significance is shown only for ZFR3 and ZFR11 with **** *P* ≤ 0.0001). (C) Fold changes in *Myd88* mRNA transcript levels were quantified relative to the YFP group (*N* = 2 biologically independent samples). Data are presented as mean + s.e.m. Statistical analysis was performed using one-way analysis of variance (ANOVA) followed by Dunnett’s multiple comparisons test. Differences were considered statistically significant at \**P* ≤ 0.05.

**Supplementary Figure 2.**
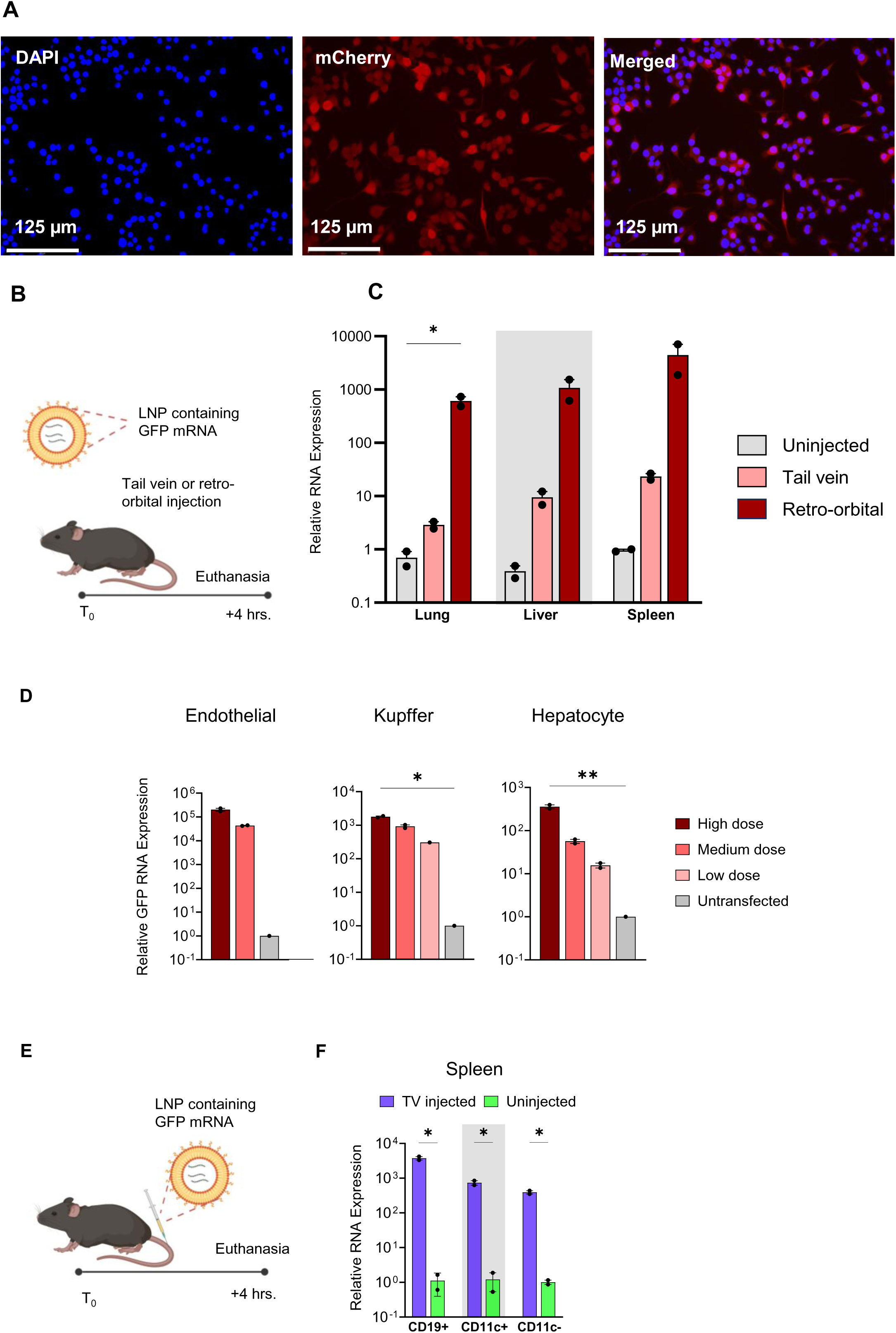
Evaluating 306O_10_ transfection efficiency and cell-type specific targeting *in vitro* and *in vivo*. (A) Fluorescence microscopy images of RAW 264.7 culture 24 hours post-transfection. Images show DAPI (nuclear staining), mCherry (anti-mCherry immunofluorescence staining), and merged channels. Scale bar, 125 μm. (B) Schematic representation of GFP-306O_10_ lipid nanoparticle delivery to C57BL/6 mice (1.2 mg mRNA/kg mouse). (C) qRT-PCR analysis of GFP expression levels in lung, liver, and spleen tissues from C57BL/6 mice four hours after systemic injection of GFP-306O_10_ via either retro-orbital (RO) or tail vein (TV) routes (N = 2 mice per group) or no injection (control, N = 2 mice). Data are normalized to uninjected controls and presented as mean + s.e.m. Statistical analysis was performed using one-way analysis of variance (ANOVA) followed by Dunnett’s multiple comparisons test. \**P* ≤ 0.05 was considered statistically significant. (D) GFP mRNA expression in primary liver cell populations (LSECs, Kupffer cells, and hepatocytes) 48 hours after GFP-306O_10_ transfection at three concentrations (high: 0.01 mg/mL, medium: 0.005 mg/mL, low: 0.0025 mg/mL). Fold changes were quantified relative to untransfected controls (N = 2 biologically independent samples). Data are presented as mean + s.e.m. Statistical analysis was performed using one-way ANOVA followed by Dunnett’s multiple comparisons test. \**P* ≤ 0.05 was considered statistically significant. (E) Schematic representation of experimental design for characterizing 306O_10_ immune cell targeting in spleen and bone marrow. C57BL/6J mice received GFP-306O_10_ (1.5 mg kg^-1^) via tail vein injection. (F) GFP mRNA expression in CD19+, CD11c+, and CD11c-populations isolated by magnetic-activated cell sorting from spleen tissues 4 hours post-injection (N = 2 mice per group). Data are presented as mean + s.e.m. Statistical analysis was performed using the unpaired parametric t-test. \**P* ≤ 0.05 was considered significant.

**Supplementary Figure 3.**
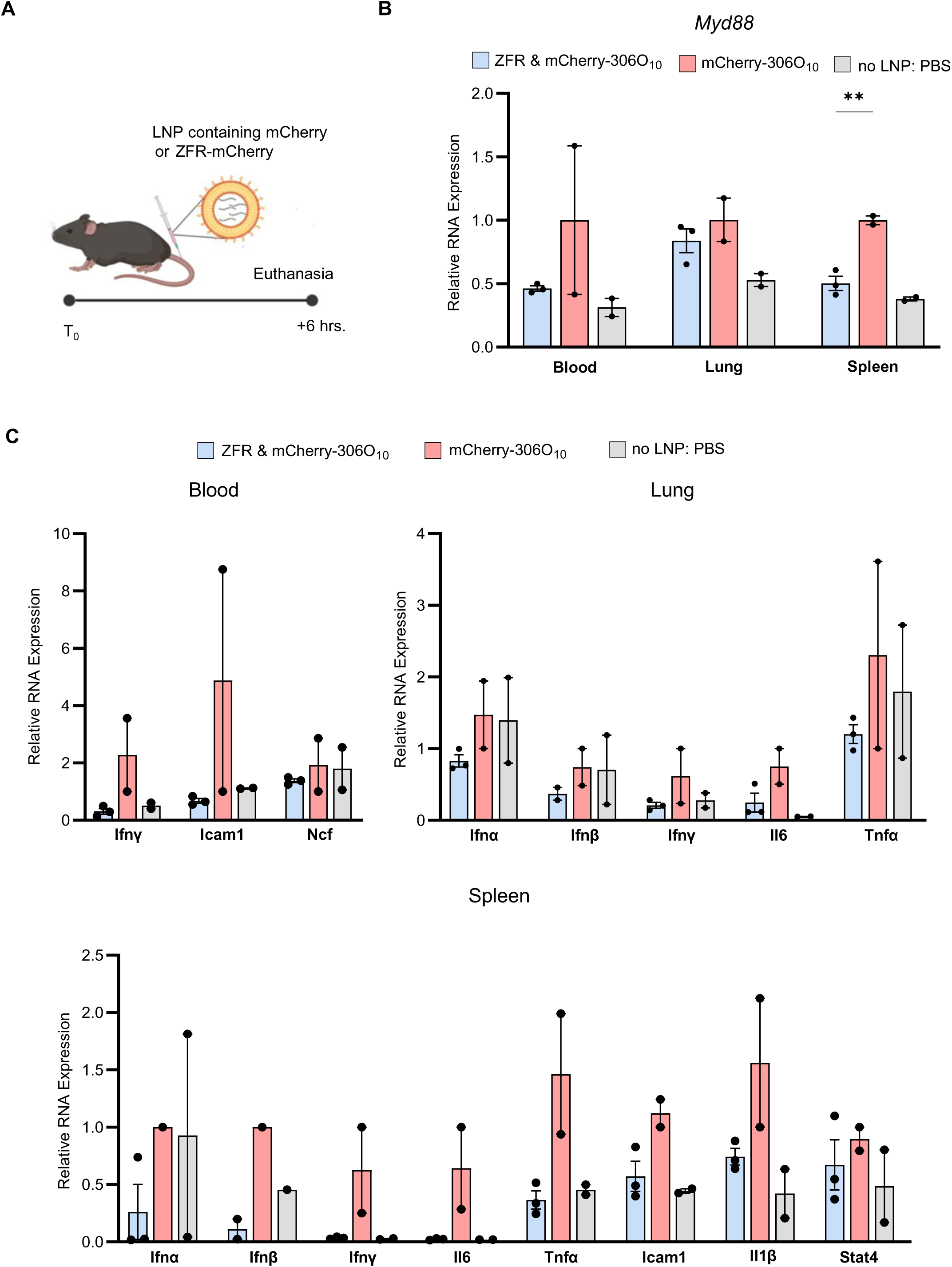
Delivery of lipid nanoparticles carrying mRNA encoding *Myd88*-targeting zinc finger enables immunomodulation under homeostatic conditions. (A) Schematic representation of experimental design. C57BL/6 mice received intravenous injection of either mCherry-306O_10_ or ZFR11-mCherry-306O_10_ (0.5 mg kg^-1^). Tissues were collected six hours post-administration for analysis. (B) qRT-PCR analysis of *Myd88* expression in blood, lung, and spleen tissues six hours after zinc finger-mediated repression (N = 3 mice for ZFR11-mCherry-306O_10_ group, N = 2 for mCherry-306O_10_ control group). Data are presented as mean + s.e.m. Statistical analysis was performed using the unpaired parametric t-test. \**P* ≤ 0.05 was considered statistically significant. (C) qRT-PCR analysis of inflammatory gene expression (*Ifn-α*, *Ifn-β*, *Ifn-γ*, *Icam-1*, *Ncf*, *Il6*, *Tnf-a*, *Il-1β*, and *Stat4*) across different tissues. Expression levels were normalized to matched tissue samples from mCherry-306O_10_-treated control mice (N = 6 mice for ZFR11-mCherry-306O_10_ group, N = 3 for mCherry-306O_10_ group). Data are presented as mean + s.e.m. Statistical analysis was performed using the unpaired parametric t-test. \**P* ≤ 0.05 was considered statistically significant.

**Supplementary Table1.**
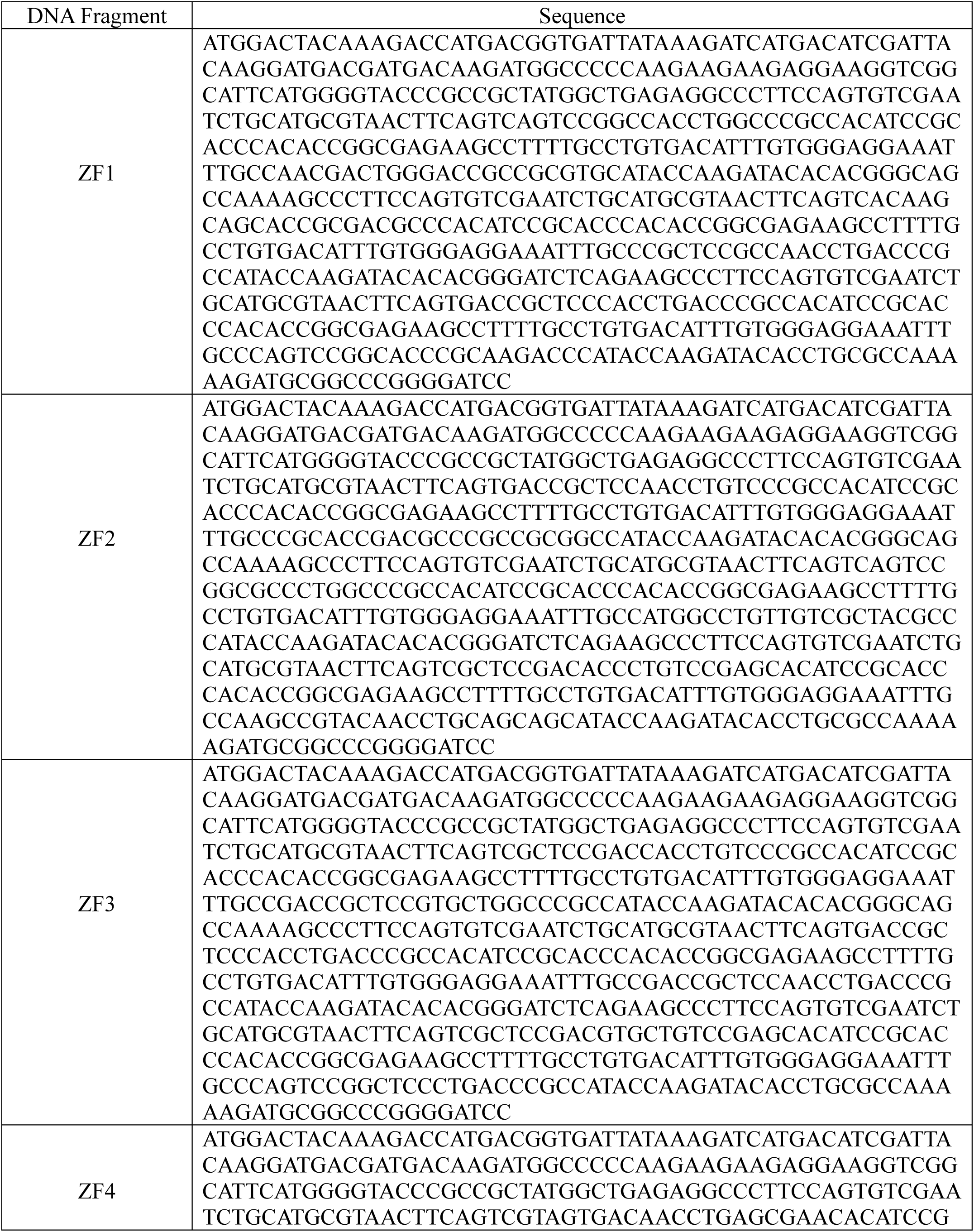
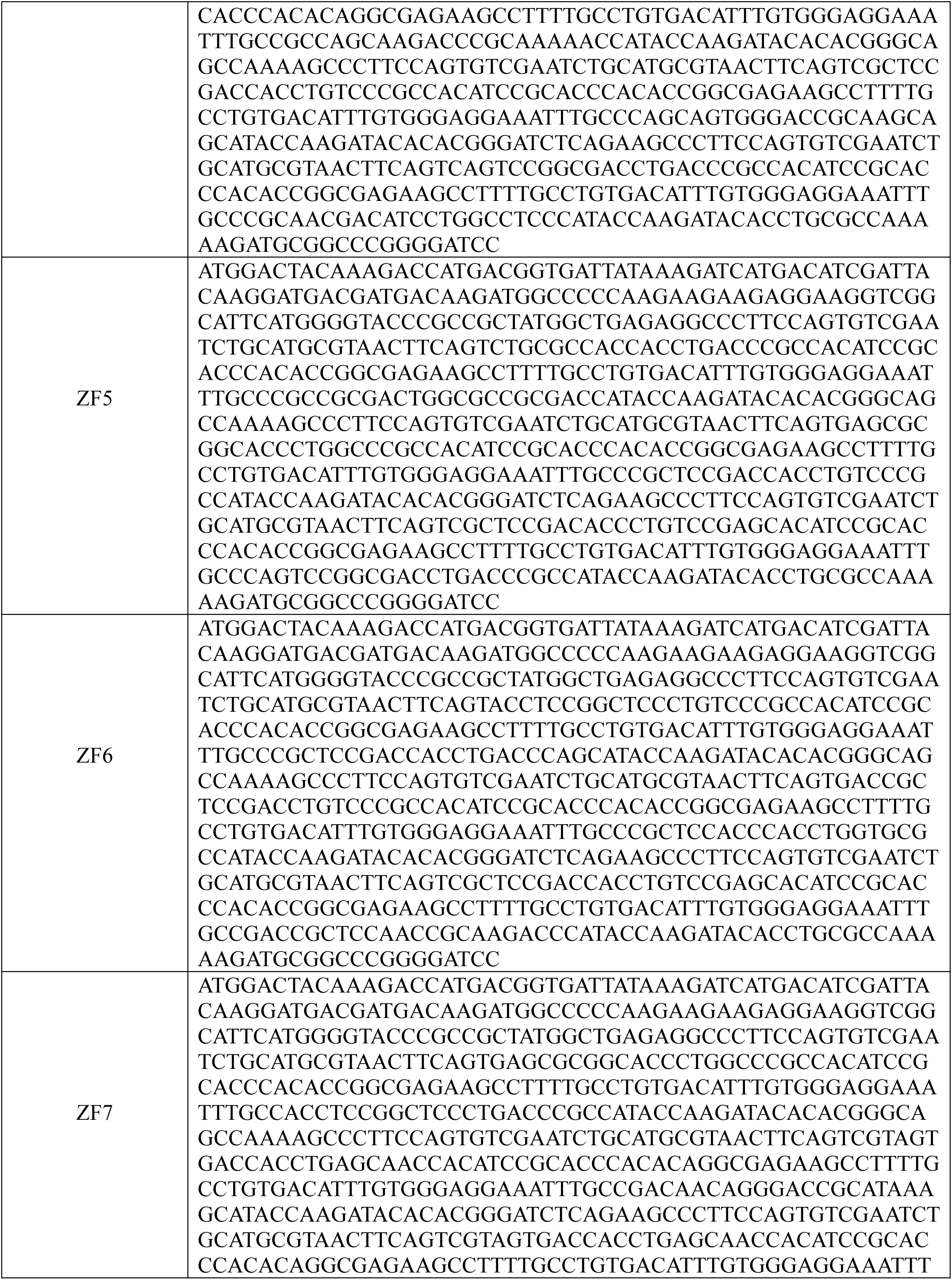
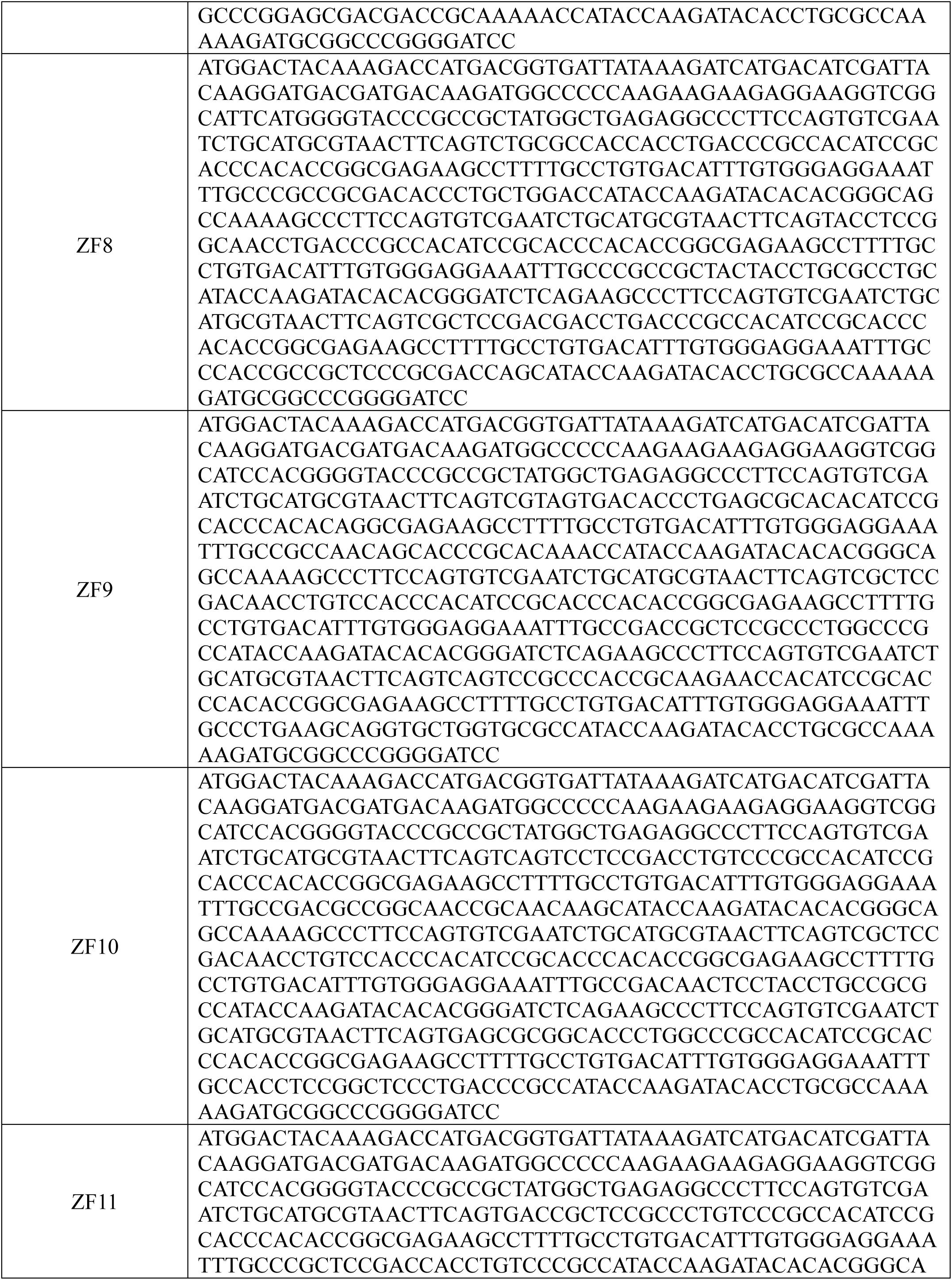
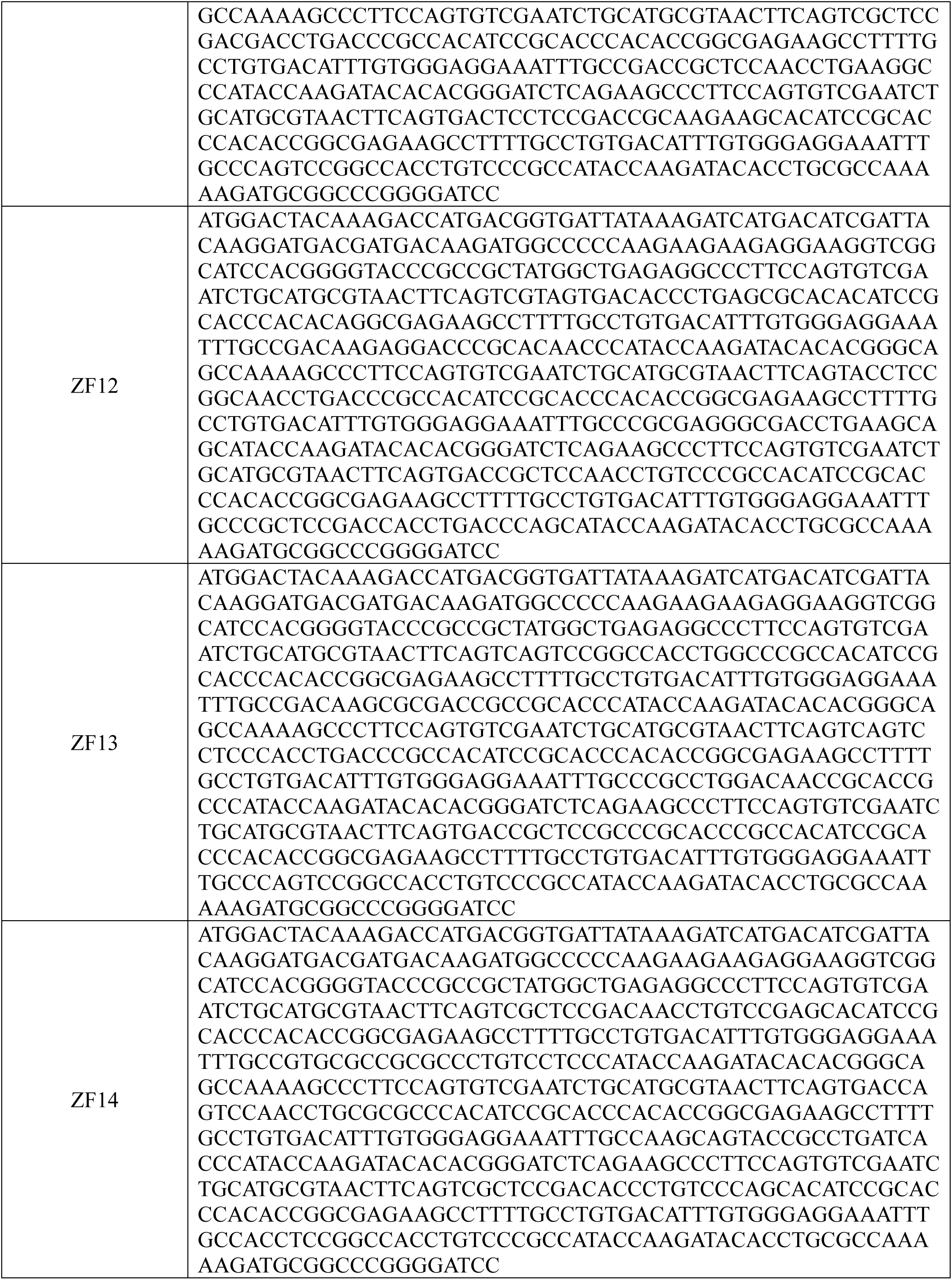
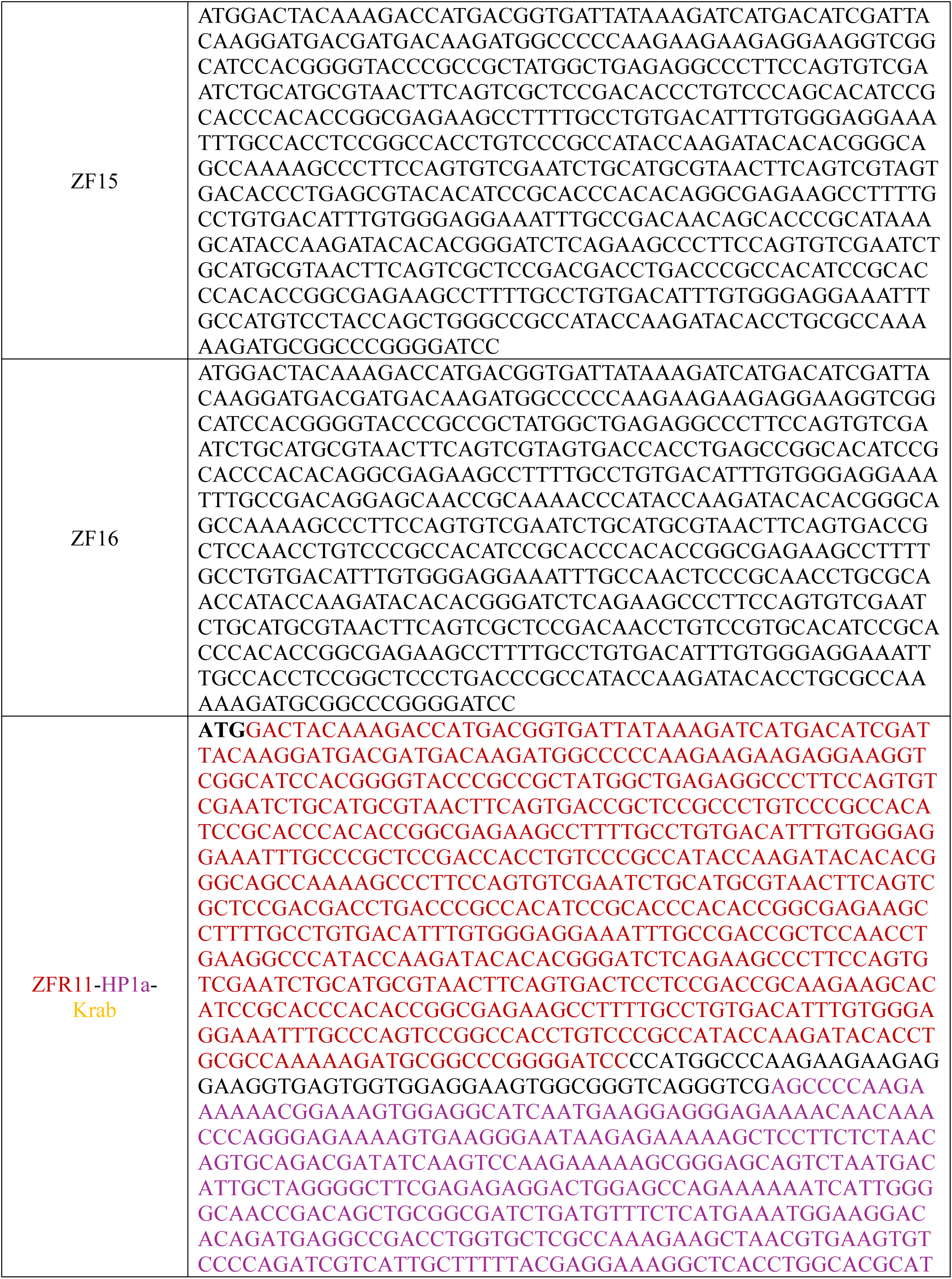
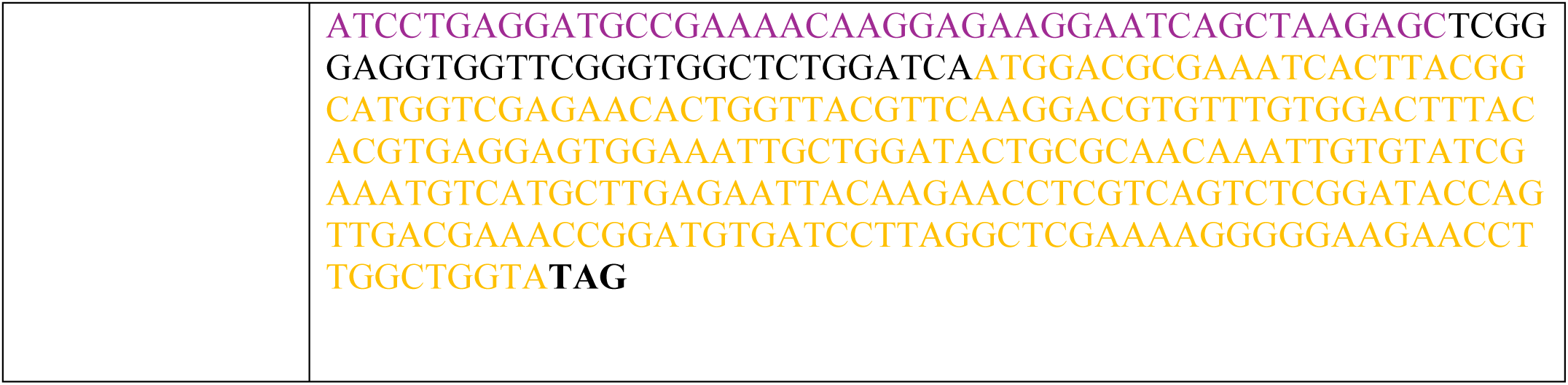
Sequences of the zinc finger DNA fragments.

**Supplementary Table 2.**
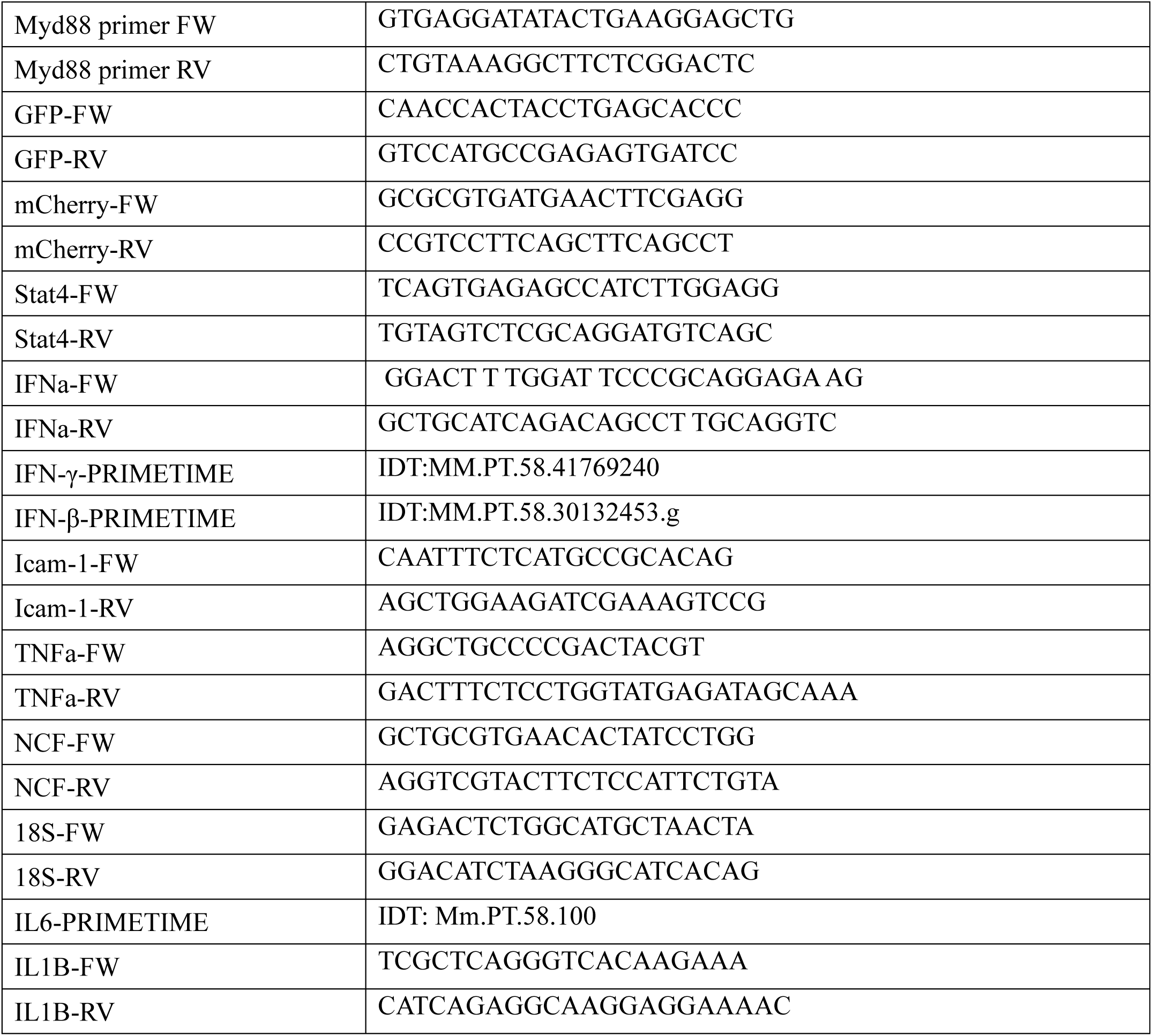
Mouse qPCR primers-5’ to 3’.

